# Targeting endothelial ERG to mitigate vascular regression and neuronal ischemia in retinopathies

**DOI:** 10.1101/2024.12.27.630529

**Authors:** Eric Ma, Christopher M. Schafer, Jun Xie, Yelyzaveta Rudenko, John T. H. Knapp, Anna M. Randi, Graeme M. Birdsey, Courtney T. Griffin

**Affiliations:** Cardiovascular Biology Research Program, Oklahoma Medical Research Foundation; Oklahoma City, USA; National Heart and Lung Institute, Imperial College London; London, UK; Department of Cell Biology, University of Oklahoma Health Sciences Center; Oklahoma City, USA

**Keywords:** Microvascular Rarefaction, Transcriptional Regulator ERG, Retina, Retinopathy of Prematurity, Diabetic Retinopathy

## Abstract

Retinopathy of prematurity (ROP) and diabetic retinopathy (DR) are ocular disorders in which a loss of retinal vasculature leads to ischemia followed by a compensatory neovascularization response. In mice, this is modeled using oxygen-induced retinopathy (OIR), whereby neonatal animals are transiently housed under hyperoxic conditions that result in central retina vessel regression and subsequent neovascularization. Using endothelial cell (EC)-specific gene deletion, we found that loss of two ETS-family transcription factors, ERG and FLI1, led to regression of OIR-induced neovascular vessels but failed to improve visual function, suggesting that relevant retinal damage occurs prior to and independently of neovascularization. Turning our attention to the initial stage of OIR, we found that hyperoxia repressed ERG expression in retinal ECs of wild type mice, raising the possibility that oxygen-induced ERG downregulation promotes vessel regression during the initiation of OIR-induced pathology. We therefore developed a murine model of EC-specific ERG overexpression and found it sufficient to prevent hyperoxia-induced vascular regression, neuronal cell death, and neovascularization in the OIR model. Importantly, ERG overexpression also improved visual function in OIR-challenged mice. Moreover, we show that both ERG and FLI1 are downregulated in the retinal vessels of human patients with early stages of DR, suggesting that neovascular disorders of the eye may share common mechanisms underlying pathological retinal capillary regression. Collectively, these data suggest that the regulation of vascular regression by EC-expressed ETS transcription factors may be adapted towards novel therapeutic approaches for the prevention and/or alleviation of ocular neovascular disorders.

## INTRODUCTION

Ischemic retinopathies are neurovascular pathologies that include two major ocular diseases: retinopathy of prematurity (ROP) in premature infants and diabetic retinopathy (DR) in the working-age population^1,2^. These diseases are characterized by an initial insult to healthy retinal vasculature, leading to capillary regression and a subsequent lack of nutrition and oxygenation (i.e., ischemia) in light-sensing neurons^1,2^. When their high nutritional demands are not met, retinal neurons generate excessive proangiogenic factors, such as vascular endothelial growth factor (VEGF), which drives compensatory angiogenesis and the formation of abnormal vascular structures known as neovascular tufts^1,2^. Current treatments suppress VEGF signaling to stabilize neovascular tufts, which can cause severe complications such as vascular leakage, hemorrhage, retinal detachment, and blindness if left untreated^3,4^. However, these treatments have important drawbacks, including variable patient efficacy, the need for repeated injections, and concerns related to broad VEGF inhibition in developing infants^1^. Moreover, by focusing on the resolution of neovascular tufts, it is unclear if these treatments address the underlying ischemic damage to retinal neurons that occurs prior to retinal neovascularization^5,6^.

There are considerable differences in the factors that drive ROP and DR. ROP is linked to the use of high oxygen incubators to support the function of underdeveloped lungs in premature infants^7^. For reasons that are still poorly understood, the increased oxygen exposure leads to vascular insufficiency via an anti-angiogenic and pro-regressive effect on retinal blood vessels^8,9^. In contrast, DR is largely restricted to adult populations and occurs as a sequela of hyperglycemia rather than hyperoxia^10^. Despite these different triggers, it is notable that both diseases follow a similar biphasic progression in which an initial phase of vascular regression causes tissue ischemia and a subsequent wave of pathological neovascularization. Therefore, the prevention of vascular regression during the initial stages of ROP and DR may offer a therapeutic window for averting the ischemic tissue damage that ultimately results in neovascularization. However, little is presently known regarding the mechanism(s) of apoptotic vascular regression or how it may be induced by hyperoxia and/or hyperglycemia during ROP and DR, respectively.

We previously studied the developmental regression of the transient ocular hyaloid blood vessel network, showing how mechanisms learned from this physiological process can be therapeutically adapted for use in a murine model of ROP^11^. This study highlighted the importance of erythroblast transformation-specific (ETS)-family transcription factors during vascular regression^11^. ETS factors play key roles in regulating various biological processes, including apoptosis, angiogenesis, cell differentiation, and proliferation^12^. In our previous work, we identified two endothelial cell (EC)-expressed ETS-family transcription factors, ETS-related gene (ERG) and Friend leukemia integration 1 (FLI1), that are downregulated in hyaloid vessels prior to regression^11^. ERG and FLI1, which share ∼65% of their protein sequences, are highly expressed by ECs, where they maintain vascular stability and quiescence^12–14^. Constitutive EC-specific deletion of *Erg* in mice leads to embryonic lethality due to severe vascular defects, and inducible deletion of *Erg* in adult ECs results in impaired physiological angiogenesis^14^. Compared to the loss of *Erg,* deletion of endothelial *Fli1* results in more subtle phenotypes, though numerous studies have shown FLI1 to have synergistic and partially compensatory functions with ERG^13,15^. As a result, the combined genetic deletion of both factors in adult ECs leads to rapid multi-organ failure and death within two weeks, primarily due to acute systemic vascular collapse characterized by hyperpermeability, hyperinflammation, and coagulopathy with spontaneous thrombosis^13^. These findings indicate that ERG and FLI1 are crucial factors regulating EC homeostasis and may play critical roles in governing vascular regression.

Here, we use murine models of inducible and EC-specific *Erg* and *Fli1* knockout and *Erg* overexpression to demonstrate how the manipulation of these ETS factors can be used either to promote or prevent vascular regression in a murine model of ROP. Using these models, we compare the impact of two distinct therapeutic strategies on retinal function: (1) promoting the regression of pathological neovascular tufts, or (2) preventing the hyperoxic regression of retinal blood vessels prior to tuft formation. Our data suggest that counteracting hyperoxia-induced ERG downregulation via EC-specific *Erg* overexpression prevents hyperoxic vessel regression, thereby avoiding ischemic damage to the retina and subsequent neovascularization. Moreover, we present evidence that ERG downregulation in retinal ECs also occurs in humans with early stages of DR, suggesting a common mechanism of pathological retinal capillary regression with ROP.

## MATERIALS AND METHODS

### Study design

This study aimed to manipulate retinal vascular regression to prevent pathologies associated with ROP. Using the murine OIR challenge model that mimics ROP progression, we first attempted to promote regression of pathological neovascular tufts with genetic deletion of the ETS transcription factors *Erg* and *Fli1* in ECs. When this failed to rescue visual function as measured by electroretinograms and optokinetic tracking in OIR-challenged mice, we next shifted focus to an earlier stage of the model in which pathological regression of normal retinal vasculature is triggered through exposure to hyperoxia. At this stage, we attempted to counteract hyperoxia-induced regression by genetically overexpressing endothelial *Erg* since we found ERG downregulation occurred in mouse retinal ECs and in cultured primary human retinal ECs after exposure to high oxygen. We also assessed ERG and FLI1 expression in the postmortem eyes of patients with non-proliferative DR to determine if these ETS factors are likewise downregulated in the early stage of this retinopathy, which is associated with pathological regression of retinal capillaries similarly to ROP.

For the mouse studies, each experimental group consisted of a minimum of six mice to ensure a robust demonstration of differences in regression phenotypes and the expression of relevant genes. The minimum animal numbers and sample sizes required to achieve statistical significance were determined by power analyses based on pilot experiments and published literature. Littermate mice were used as controls for all the animal studies. The investigators were blinded to the sample analyses and quantification. Sample sizes and specific statistical tests used for each experiment are outlined in the figure legends.

### Study approvals

Histological sections from deidentified donor eyes from patients with non-proliferative DR (NPDR) and from nondiabetic patients were obtained from Lions Gift of Sight Eye Bank (Saint Paul, MN). All methods were performed in accordance with federal and institutional guidelines. Protocols involving mouse breeding, tamoxifen induction, the OIR model, and visual function testing followed the guidelines of the Association for Research in Vision and Ophthalmology (ARVO) Statement for the Use of Animals in Ophthalmic and Visual Research. All studies were conducted following protocols approved by the Oklahoma Medical Research Foundation Institutional Animal Care and Use Committee (#23-12 and #24-27).

### Animals

Mice used in this study included C57BL/6J (The Jackson Laboratory; #000664), *Erg^flox^* (gift of Joshua Wythe, Baylor College of Medicine; available through The Jackson Laboratory; #030988)^16^, *Fli1^flox^* (gift of Maria Trojanowska, Boston University)^17^, and *Cdh5(PAC)-Cre^ERT2^*(gift of Ralf Adams, Max Planck Institute for Molecular Biomedicine; available through Taconic; #13073)^18^. *ROSA^Erg/Erg^* (generated by Graeme Birdsey and Anna Randi, Imperial College London), were constructed by genOway (Lyon, France). Briefly, a knock-in vector was designed, containing murine *Erg* cDNA inserted upstream of an IRES-tdTomato-Reporter cassette. A loxP flanked Stop cassette was inserted between a CAG promoter and the *Erg* cDNA to allow expression to be dependent upon induction by Cre recombinase. The inducible targeting vector was inserted into the permissive *Rosa26* locus of C57BL/6 mice. Mice were housed at the Oklahoma Medical Research Foundation animal facility in a specific pathogen-free (SPF) environment at 25°C under a 12/12 hr light/dark cycle. Both male and female were included in all studies. Genotyping of *ROSA^Erg/Erg^* was performed using the primers 5′-GTTTTGGAGGCAGGAAGCACTTGC-3′, 5′-GCAGTGAGAAGAGTACCACCATGAGTCC-3′, and 5′-CGAGGCGGATCACAAGCAATA-3′. Genotyping of *Erg^flox^* was performed using the primers 5′-GAGATGGCGCAACGCAATTAATG-3′, 5′-AGAGTCTCTGCACACAGAACTTCC-3′, and 5′-AATGCTCTGGTAAGGCACACAAGG-3′, as previously described^11^. Genotyping of *Fli1^flox^* was performed using the primers 5′-GACTCAAACCAGGGAAAGTTGC-3′, and 5′-TTGGGAAGGTGGAATCTAGCAG-3′. Genotyping of *Cdh5(PAC)-Cre^ERT2^* mice was performed using the primers 5′-TCCTGATGGTGCCTATCCTC-3′ and 5′-CGAACCTGGTCGAAATCAGT-3′. Tamoxifen-induced *Cdh5(PAC)-Cre^ERT2^* activation was achieved by oral gavage of 20 mg/ml tamoxifen (Cat#T5648, Sigma) dissolved in peanut oil, administered to pups at P5-P7 (3 µl/day) or P16-P17 (5 µl/day). An eyedrop route of tamoxifen administration (20 mg/ml, 3 µl/day from P5 to P7) was used for *Erg^iECoe^* and littermate control mice that were subsequently used for electroretinogram measurements at P21.

### Oxygen-induced Retinopathy (OIR) model

The mouse OIR model was performed as described previously^19^. Neonatal mouse pups were housed in room air (RA; 21% oxygen) until P7. Then, mouse pups were housed in hyperoxia chambers (75% oxygen; Biospherix) for 5 days from P7 to P12, during which time hyperoxia-induced retinal vascular regression occurred. Mice were next removed from the hyperoxia chambers and were housed in RA until terminal experiments were performed. Note that the peak time of neovascular tuft formation occurred at P17. Visual function tests (OKT and electroretinograms) were performed at P21. Littermate *Cre^ERT2^*-negative mice were used as controls in the OIR model and received tamoxifen treatment at the same time as their experimental counterparts. Age-matched mice were used as RA controls for animals that were subjected to the OIR model. Both sexes were used for experiments.

### Immunofluorescent staining and imaging

Human eyes were collected within 32 hours postmortem and were overnight fixed in Davidson’s fixative. Antigen epitopes were retrieved with heat induction using EnVision FLEX Target Retrieval Solution (Citrate buffer pH 6.1, Agilent Technologies, Inc., Santa Clara, CA). Tissue autofluorescence was quenched with 0.1% Sudan Black. Mouse whole-mount retinal staining was performed as described previously^20^. Briefly, mouse eyeballs were enucleated and fixed in 4% paraformaldehyde (PFA) in phosphate-buffered saline (PBS, pH 7.4) for 1 hr. Retinas were isolated from fixed eyeballs under a dissection microscope. The retinas were incubated in a blocking buffer [3% bovine serum albumin (BSA), 10% donkey serum, and 0.5% Triton X-100 in PBS] overnight at 4°C. The retinas were incubated in primary antibodies overnight at 4°C followed by secondary antibodies overnight at 4°C and were then flatmounted. 4’,6-diamidino-2-phenylindole (DAPI) was used for counterstaining, and ProLong Diamond Antifade Mountant (Cat P36930, Invitrogen) was used as a mounting buffer. Images were collected using a Nikon AXR confocal microscope and analyzed using Nikon Elements software. SWIFT_NV plugin in ImageJ software was used to quantify neovascular and avascular areas in the retinal flatmounts^21^. Details of primary and secondary antibodies used in this study are listed in Supplementary Table 1.

### Terminal Deoxynucleotidyl Transferase-dUTP Nick End Labeling (TUNEL) staining

Ocular cryosections were prepared as described previously^20^. Sagittal sections (5 µm) were generated through the peripheral and central regions of the retina and through the optic nerve. TUNEL staining was performed using In Situ Cell Death Detection Kit (Cat 11684795910, Roche) according to the manufacturer’s protocol^22^. Sections were incubated in TUNEL reaction mixture for 1 hr at 37°C. DAPI was used for counterstaining. TUNEL^+^ cells were quantified using ImageJ software.

### Immunoblot analysis

Immunoblot analysis was performed as described previously^23^. Briefly, retinas were dissected and snap-frozen in liquid nitrogen and stored at -80°C until use. Tissues were lysed in RIPA buffer (Sigma) supplemented with 1% proteinase inhibitor cocktail (Sigma). After sonication, the samples were centrifuged at 12,000 x g for 10 min at 4°C. Protein concentrations within the supernatant were determined by BCA assay (ThermoFisher). Samples with equal amounts of proteins were resolved in SDS-PAGE gels (Invitrogen) and transferred to nitrocellulose membranes (Invitrogen). After blocking with 10% nonfat milk in Tris-buffered saline with 0.1% Tween 20 (TBST), the membranes were incubated with primary antibodies overnight at 4°C, followed by incubation with secondary antibodies for 2 hr. All antibodies were diluted in TBST containing 5% BSA. Details of primary and secondary antibodies used in this study are listed in Supplementary Table 1.

### Quantitative reverse transcription polymerase chain reaction (qPCR)

Cultured endothelial cells were lysed in Trizol reagent (Invitrogen). RNA was extracted using an RNeasy Mini Prep Kit (QIAGEN). cDNA was generated using an iScript cDNA Synthesis Kit (Bio-Rad). Target gene mRNA levels were measured by qPCR using SsoAdvanced Universal SYBR Green Sypermix (BioRad) and analysis on a CFX96 Real-Time PCR thermocycler (Bio-Rad). The following primers were used for measuring human *ERG* mRNA: 5′-AACGAGCGCAGAGTTATCGTGC-3′ and 5′-GTGAGCCTCTGGAAGTCGTCC-3′. Human *18s* and *ACTB* were used as housekeeping gene transcripts for normalization and were measured using the primers 5′-CCCGAAGCGTTTACTTTGAAA-3′ and 5′-CGCGGTCCTATTCCATTATTC-3′ (for *18s*) and the primers 5′-CTCTTCCAGCCTTCCTTCCT-3′ and 5′-AGCACTGTGTTGGCGTACAG-3′ (for *ACTB*).

### Electroretinograms

Scotopic and photopic responses to electroretinograms were measured using a Celeris system (Diagnosys), as described previously^24,25^. Briefly, mice were dark-adapted overnight (≥ 18 hr). Under dim red light, mice were anesthetized by an intraperitoneal injection of 120 µg/g ketamine and 9 µg/g xylazine diluted in saline. Pupils were sequentially dilated with 0.5% proparacaine hydrochloride ophthalmic solution (Sandoz), 1% tropicamide hydrochloride ophthalmic solution (Sandoz), and 10% phenylephrine hydrochloride ophthalmic solution (Sandoz). Body temperature was maintained at 37°C on the Celeris platform. Systane lubricant eye gel (Alcon) was applied to conduct the electrical signal between the cornea and electrode. The contralateral unstimulated eye was used as the reference. Scotopic electroretinograms were performed using light flashes of intensities ranging from -2.0 to 1 log cd.s/m^2^. For each intensity, at sufficient intervals (60 to 300s), mice were allowed to readapt to darkness, as to recover from any photobleaching effects. Photopic electroretinogram recordings were performed after 5-minute intervals of white light adaptation at 10 cd.s/m^2^ to desensitize rods. For photopic electroretinograms, the cone response was measured at 3 different light intensities (1, 3, and 10 cd.s/m^2^) in the presence of white light (10 cd.s/m^2^).

### Optokinetic Tracking (OKT)

Visual acuity thresholds were measured using an Optometry apparatus and software (Cerebral Mechanics) as described previously^26^. Briefly, the spatial frequency and contrast sensitivity were assessed in awake, freely moving mice at P21. Mice were placed on a pedestal inside a chamber with a virtual cylinder consisting of vertical lines projected on 4 computer screens. The vertical lines rotated at varying frequencies, and tracking behavior was assessed in a stepwise manner. Spatial frequency was represented as the highest frequency (cycle/degree) at which mice tracked the rotating cylinder. The contrast threshold was measured at a spatial frequency of 0.064 cycles/degree, based on a previous publication of the OIR model^27^. Contrast sensitivity was identified as the highest value that still elicited a response in the mouse. Both spatial frequency and contrast sensitivity were measured from 3 different litters at P21.

### Cell culture

Primary human retinal microvascular endothelial cells (HREC; ACBRI 181, Cell Systems) or human umbilical vein endothelial cells (HUVECs; PCS-100-010, ATCC) were cultured in endothelial complete media (4Z0-500, Cell Systems) or EGM2 (Lonza), respectively. Only cells that were passed from 2 to 8 times were used for experiments. Culture under variable oxygen levels was accomplished using a ProOx model C21 (BioSpherix, NY). For siRNA treatments, 1.0-1.5X10^5^ HUVECs were plated in 6-well dishes and incubated overnight at 37°C. The following morning, media was replaced with OptiMEM (Thermo) containing either Silencer Select Negative Control No.1 siRNA or *hTRIM25*-targeting siRNA (Thermo; s15206) diluted in Lipofectamine RNAiMAX transfection agent (Thermo) following the manufacturer’s guidelines.

### Statistics

At least six mice per group were used for all animal experiments, and mice were derived from at least two separate litters. At least three technical replicates were performed for in vitro experiments. Results are presented as mean ± SD. Tests for normality (D’Agostino-Pearson and Shapiro-Wilk) and equal variance (Brown-Forsythe) were conducted to determine the appropriate parametric or nonparametric statistical models. Statistical analyses were conducted using a two-tailed Student’s *t* test for comparison of two groups, or using a one-way ANOVA followed by the Student-Newman-Keuls multiple comparison test when more than two groups were compared. *P* values of <0.05 were considered statistically significant. Prism 10.0 software (GraphPad) was used for all statistical assessments.

## RESULTS

### Deletion of endothelial ERG and FLI1 promotes neovascular tuft regression

Oxygen-induced retinopathy (OIR) is a murine model of ROP in which neonatal mice are housed under hyperoxia (75% O2) from postnatal day (P) 7 to P12, leading to oxygen-induced regression of retinal blood vessels (Fig. 1A). As in humans, the resulting vascular insufficiency leads to a compensatory phase of retinal neovascular tuft formation that peaks around P17^19^. Previously, we showed that neovascular tufts can be selectively regressed by treatment with a pharmacological inhibitor of ETS family transcription factors^11^. However, the specific identities of the relevant ETS factors, which include 28 genes in humans and 27 genes in mice, was not determined.

**Figure 1.**
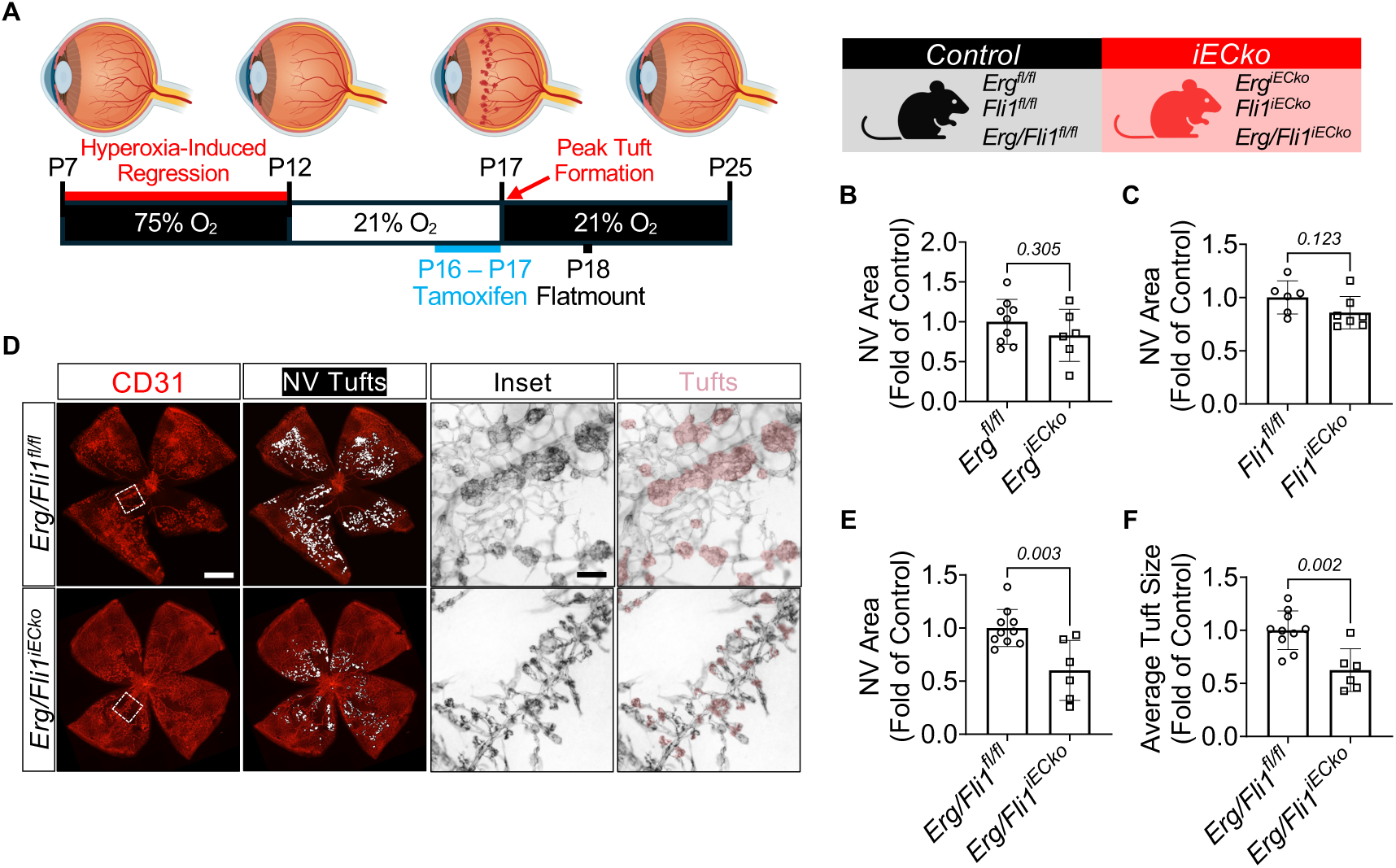
EC-specific deletion of both *Erg* and *Fli1* (*Erg/Fli1^iECko^*) promotes NV tuft regression. **(A)** Schematic of subjecting single knockout (KO) and littermate flox control mice (*Erg^iECko^* vs. *Erg^fl/fl^*; *Fli1^iECko^*vs. *Fli1^fl/fl^*) and double KO and littermate flox control mice (*Erg/Fli1^iECko^* vs. *Erg/Fli1^fl/fl^*) to the oxygen-induced retinopathy (OIR) model. Pups were housed in room air (21% O_2_) from birth to P7 and were then exposed to hyperoxia (75% O_2_) for 5 days from P7 to P12, when pathological retinal vascular regression occurs. Pups were returned to room air from P12 to P18, and P17 was the time of peak neovascular (NV) tuft formation. To induce *Cdh5(PAC)-Cre^ERT2^* activation, tamoxifen was administered orally at P16 and P17. Retinas were harvested at P18 for flatmount staining of CD31^+^ retinal vessels. (**B, C**) Quantification of NV areas from *Erg^iECko^* mice and controls (n = 6-9 mice) and from *Fli1^iECko^* mice and controls (n = 6-7 mice). (**D**) Representative images of CD31-stained retinal flatmounts from P18 *Erg/Fli1^iECko^* and littermate control mice. The ImageJ plugin SWIFT_NV^21^ was used to label and quantify NV tuft areas (white in the whole flatmounts). The inset monochrome images correspond to the white squares in the images at the far left; tufts are pseudocolored in pink. Scale bar: 500 µm in whole flatmounts and 50 µm in the magnified inset images. (**E**) Quantification of NV areas from *Erg/Fli1^iECko^* and control mice (n = 6-10 mice). (**F**) Quantification of average tuft sizes from *Erg/Fli1^iECko^* and control mice (n = 6-10 mice). Data are presented as mean ± SD. *Erg^iECko^*: *Erg^flox/flox^;Cdh5(PAC)-Cre^ERT2^*, *Fli1^iECko^*: *Fli1^flox/flox^;Cdh5(PAC)-Cre^ERT2^*, NV: Neovascular, OIR: Oxygen-induced retinopathy.

Because ERG and FLI1 are EC-expressed ETS family transcription factors that play a critical role in maintaining vascular homeostasis^12,13^, we hypothesized that inhibition of one or both of these factors would lead to the regression of OIR-induced neovascular tufts. We therefore generated *Erg^flox/flox^;Cdh5(PAC)-Cre^ERT2^*(*Erg^iECko^*) and *Fli1^flox/flox^;Cdh5(PAC)-Cre^ERT2^*(*Fli1^iECko^*) mice to enable tamoxifen-inducible and EC-specific deletion of *Erg* and *Fli1*, respectively. *Erg^iECko^* and *Fli1^iECko^* mice were exposed to 75% O_2_ from P7 to P12, and tamoxifen was administered at P16 and P17 after neovascular tufts had already formed (Fig. 1A). At P18, retinal flat mount preparations were analyzed by immunofluorescence for the EC marker CD31, and the ImageJ plugin SWIFT_NV^21^ was used to quantify total neovascular area, neovascular tuft number, and avascular area (Fig. 1B, 1C, and Suppl Fig. 1A-C).

Though there was a trend towards less neovascularization for both *Erg^iECko^* and *Fli1^iECko^* mice, neither genotype reached significance (Fig. 1B, 1C). ERG and FLI1 have highly homologous DNA binding domains^28^ and play synergistic roles in maintaining endothelial homeostasis^15^, suggesting that the presence of either protein may functionally compensate for the loss of the other. We therefore generated *Erg^flox/flox^Fli1^flox/flox^;Cdh5(PAC)-Cre^ERT2^*(*Erg/Fli1^iECko^*) mice to achieve EC-specific deletion of both *Erg* and *Fli1* genes and subjected them to the OIR model, as described above (Suppl Fig. 1D, 1E). In contrast to the individual knockouts, simultaneous deletion of *Erg* and *Fli1* had an additive effect, resulting in a significant reduction (∼40%) in neovascular area compared to littermate controls (Fig. 1D, 1E). Moreover, average tuft sizes were significantly reduced in *Erg/Fli1^iECko^* mice (Fig. 1D, 1F), and this was likely due to tuft regression rather than compromised tuft growth since there were no significant differences in overall tuft numbers between control and mutant genotypes (Suppl Fig. 1C). Meanwhile, for mice raised in room air, there were a comparable number of empty basement membrane sleeves (CD31^-^ColIV^+^) between *Erg/Fli1^iECko^* mice and littermate controls (Suppl Fig. 1F, 1G). Since empty basement membrane sleeves are indicative of capillary regression^29^, these data suggest that regression occurs specifically in tufts of *Erg/Fli1^iECko^* mice but not in the normal retinal vasculature.

### Visual function is not improved following *Erg/Fli1^iECko^*-induced neovascular tuft regression

Our success in regressing neovascular tufts in *Erg/Fli1^iECko^* mice prompted us to ask if this led to an improvement in OIR-induced visual defects^30,31^. We therefore exposed *Erg/Fli1^iECko^* mice to 75% O_2_ from P7 to P12, induced gene deletion as before at P16/P17, and measured retinal function and visual acuity at P21 using electroretinogram and optokinetic tracking (OKT), respectively (Fig. 2A). Using both scotopic and photopic electroretinograms, we observed similar decreases in a-and b-wave amplitudes for OIR *Erg/Fli1^iECko^* and OIR control mice compared to mice of both genotypes that were raised in room air (Fig. 2B-2F). Similarly, both genotypes had comparable reductions in their threshold for spatial frequency detection as determined by OKT (Fig. 2G). Therefore, despite the regression of neovascular tufts that we achieved by deleting both *Erg* and *Fli1* from ECs (Fig. 1D-F), we observed no signs of improved visual function in OIR-challenged *Erg/Fli1^iECko^*mice.

**Figure 2.**
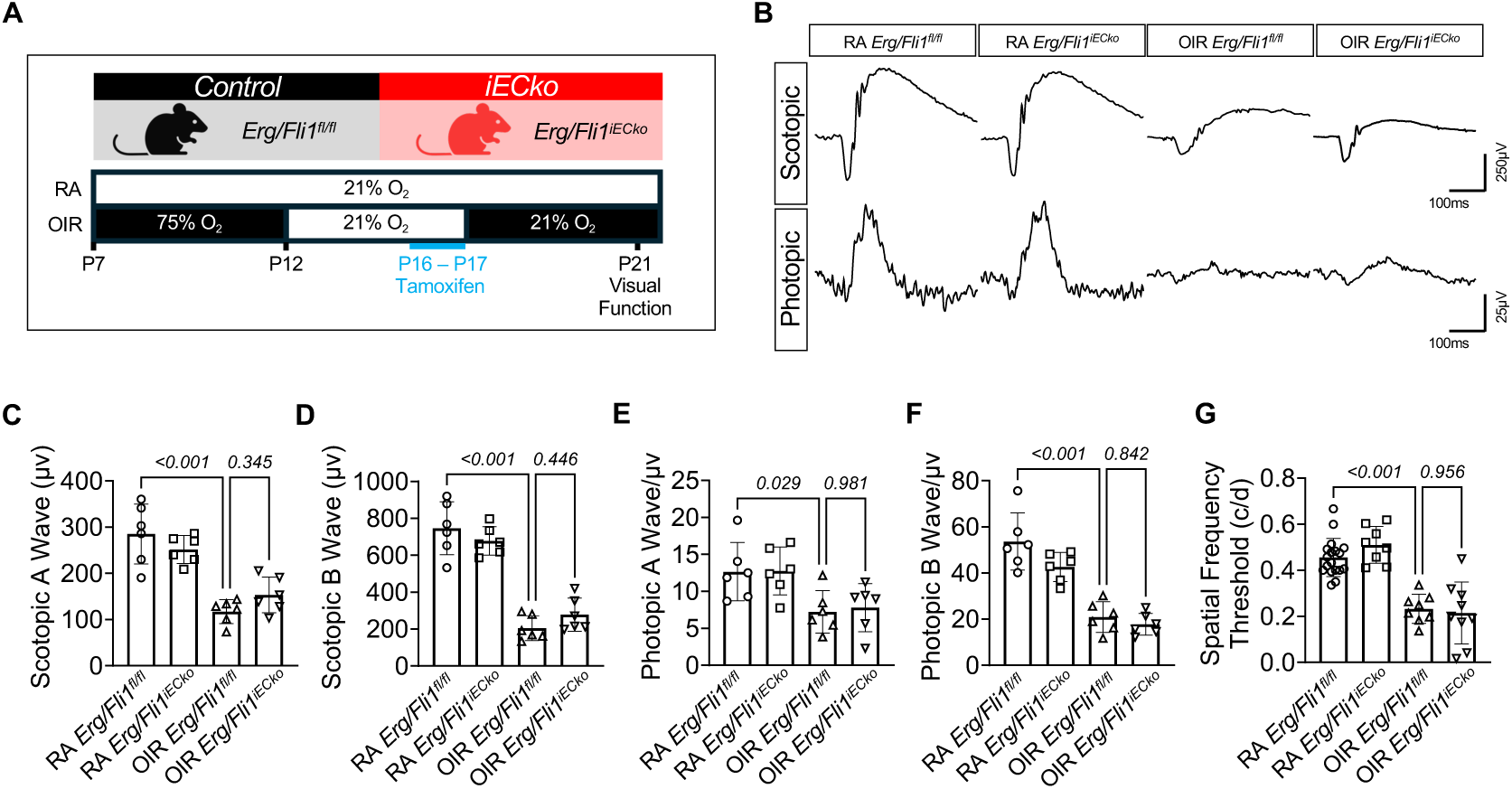
NV tuft regression in *Erg/Fli1^iECko^* mice does not improve OIR-induced visual defects. **(A)** Schematic of testing the effects of the OIR model on visual function in *Erg/Fli1^iECko^*and littermate control (*Erg/Fli1^fl/fl^*) mice. Note that age-matched mice of both genotypes that were raised in room air (RA) served as negative controls for the OIR model. Tamoxifen was administered orally to mice of both genotypes at P16 and P17 to induce Cre activation. Visual function was measured by electroretinogram and optokinetic tracking (OKT) at P21. (**B**) Representative curves of scotopic and photopic amplitudes from the four different groups. Scale bar labels are shown at the bottom right. (**C-F**) Quantification of scotopic a-wave (C), scotopic b-wave (D), photopic a-wave (E), and photopic b-wave (F) readings from the four different groups (n = 6 mice). (**G**) Quantification of spatial frequency threshold (c/d) measured by OKT (n = 8-16 mice). Data are presented as mean ± SD. OIR: Oxygen-induced retinopathy, OKT: Optokinetic tracking, RA: Room air.

In wild type mice subjected to OIR, revascularization of the central retina occurs spontaneously following the resolution of neovascular tufts by P25^32^. We reasoned that compromised revascularization might contribute to the visual defects we detected in *Erg/Fli1^iECko^*mice at P21. However, we observed persistent visual defects in OIR-treated wild type mice even after the retina had been fully revascularized at P30 (Suppl Fig. 2), suggesting that visual defects associated with the OIR model are likely due to neuronal damage that occurred prior to the neovascular phase rather than to failed revascularization of the retina after tuft resolution.

### Endothelial ERG expression is downregulated by hyperoxia exposure during OIR

Like ROP, OIR-induced neovascularization occurs because of an initial hyperoxic challenge that leads to vascular regression and the impairment of nutrient delivery to the metabolically consumptive retina. In our hands, hyperoxia-induced regression of central retinal vessels was rapid in the OIR model, with a loss of nearly one third of all retinal vessels by 24hr of hyperoxia exposure from P7 to P8 in wild type mice (Fig. 3A, 3B). Following this, vascular regression spread outward from the central retina during the subsequent 4 days of hyperoxia exposure from P8 to P12. Loss of endothelial ERG during this phase of hyperoxia exposure has previously been reported^33^, and our discovery that neovascular tuft regression is accelerated in *Erg/Fli1^iECko^*mice led us to ask if the downregulation of either transcription factor might also contribute to hyperoxia-induced vessel regression. We therefore assessed endothelial ERG and FLI1 expression in the initial 24hr window of hyperoxia exposure to determine if they were downregulated during this period in which the most substantial amount of retinal vascular regression occurred.

**Figure 3.**
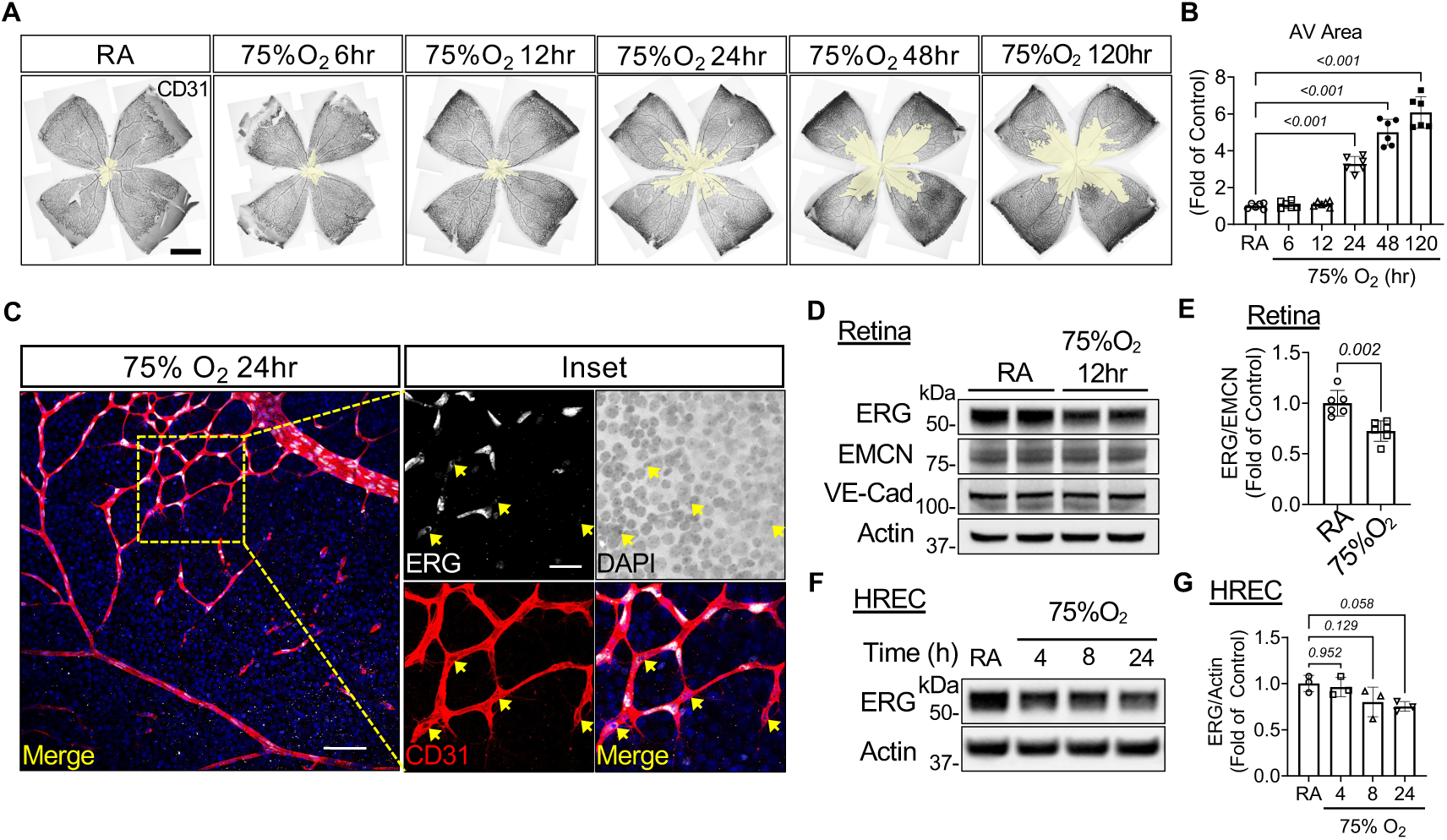
Hyperoxia downregulates ERG in wild type retinal ECs in vivo and in vitro. (**A**) Representative images of CD31-stained retinal flatmounts from wild type C57Bl/6J pups exposed to 75% O_2_ for the indicated times. Yellow layer indicates avascular (AV) area. Scale bar: 500 µm. (**B**) Quantification of AV areas in (A) (n = 6 mice). (**C**) Representative immunostaining of ERG, CD31, and DAPI in retinas from P8 wild type C57Bl/6J pups after exposure to 75% O_2_ for 24 hr. Yellow arrows indicate EC nuclei with low ERG signal at the capillary regression front. Scale bar: 100 μm in the merged image and 20 μm in the inset. (**D**) Representative immunoblots from whole retinas of P7 littermate wild type pups exposed to room air (RA) or to 75% O_2_ for 12 hr. (**E**) Densitometry quantification of ERG normalized to endomucin (EMCN) (n = 6 mice). (**F**) Representative immunoblots from primary human retinal microvascular endothelial cells (HREC) cultured under RA for 24 hr or 75% O_2_ for the indicated time. (**G**) Densitometry quantification of normalized ERG in (F) (n = 3 technical replicates). Data are presented as mean ± SD. AV: Avascular Area, EMCN: Endomucin, HREC: Human retinal microvascular endothelial cells, RA: Room air, VE-Cad: Vascular endothelial cadherin.

Immunofluorescence imaging for ERG at 24hr exposure to 75% O_2_ revealed robust nuclear expression in most retinal ECs (Fig. 3C). However, nuclear ERG was notably absent from a subset of ECs (Fig. 3C, yellow arrows) located along the border of the retinal avascular region where active vessel regression was occurring. Furthermore, immunoblot analysis of whole retina tissue lysates revealed a reduction in ERG expression in wild type mice exposed to hyperoxia for 12hr compared to littermates exposed to room air (Fig. 3D, 3E). Importantly, ERG expression was normalized to the capillary marker endomucin (EMCN; Fig. 3D, 3E), indicating that the observed reduction in ERG expression occurred in viable ECs and was not due to the overall loss of retinal vessels. By contrast, a similar reduction in retinal FLI1 expression was not observed after mice were exposed to hyperoxia for 12hr (Suppl Fig. 3A, 3B).

To determine if oxygen may have a direct effect on ERG expression in ECs, we cultured primary human retinal endothelial cells (HRECs) under 21% and 75% O_2_ and assessed ERG expression by immunoblot. By 8 and 24hr of 75% O2, we observed a moderate reduction of ERG expression (Fig. 3F, 3G). Like our in vivo findings, oxygen had little effect on FLI1 expression in vitro (Suppl Fig. 3C, 3D). We and others have previously reported that ERG downregulation can occur after its ubiquitination and subsequent proteasomal degradation, and TRIM25 is a ubiquitin ligase that putatively marks ERG for proteolysis in certain contexts^34,35^. Intriguingly, we found that oxygen exposure upregulates TRIM25 expression in HRECs, suggesting a potential mechanism for O_2_-dependent ERG downregulation (Suppl Fig. 3E). In agreement with this, siRNA-mediated deletion of *TRIM25* in ECs led to a significant upregulation of ERG protein without affecting *ERG* mRNA transcript expression (Suppl Fig. 3F-3H). Collectively, our in vivo and in vitro data indicate that the expression of ERG is more robustly regulated by oxygen exposure in retinal ECs than is FLI1 expression, with higher oxygen levels resulting in more downregulation of ERG.

### Overexpression of endothelial ERG reduces hyperoxia-induced vascular regression and neovascularization

Since we found that endothelial ERG downregulation correlated with hyperoxia-induced vascular regression, we hypothesized that the overexpression of endothelial ERG would counteract this process. We therefore developed a mouse in which murine *Erg* coding sequence was inserted into the ROSA26 locus downstream of a floxed STOP cassette. These mice were crossed with the EC-specific *Cdh5(PAC)-Cre^ERT2^*line^18^. Administration of tamoxifen to *ROSA^Erg/Erg^;Cdh5(PAC)-Cre^ERT2^* (*Erg^iECoe^*) mice from P5 to P7 led to a ∼7-fold upregulation of ERG protein in whole retinas that could be detected by immunoblot and immunofluorescence at P8 compared to littermate controls (*ROSA^Erg/Erg^*) subjected to the same tamoxifen administration scheme (Fig. 4A-D).

**Figure 4.**
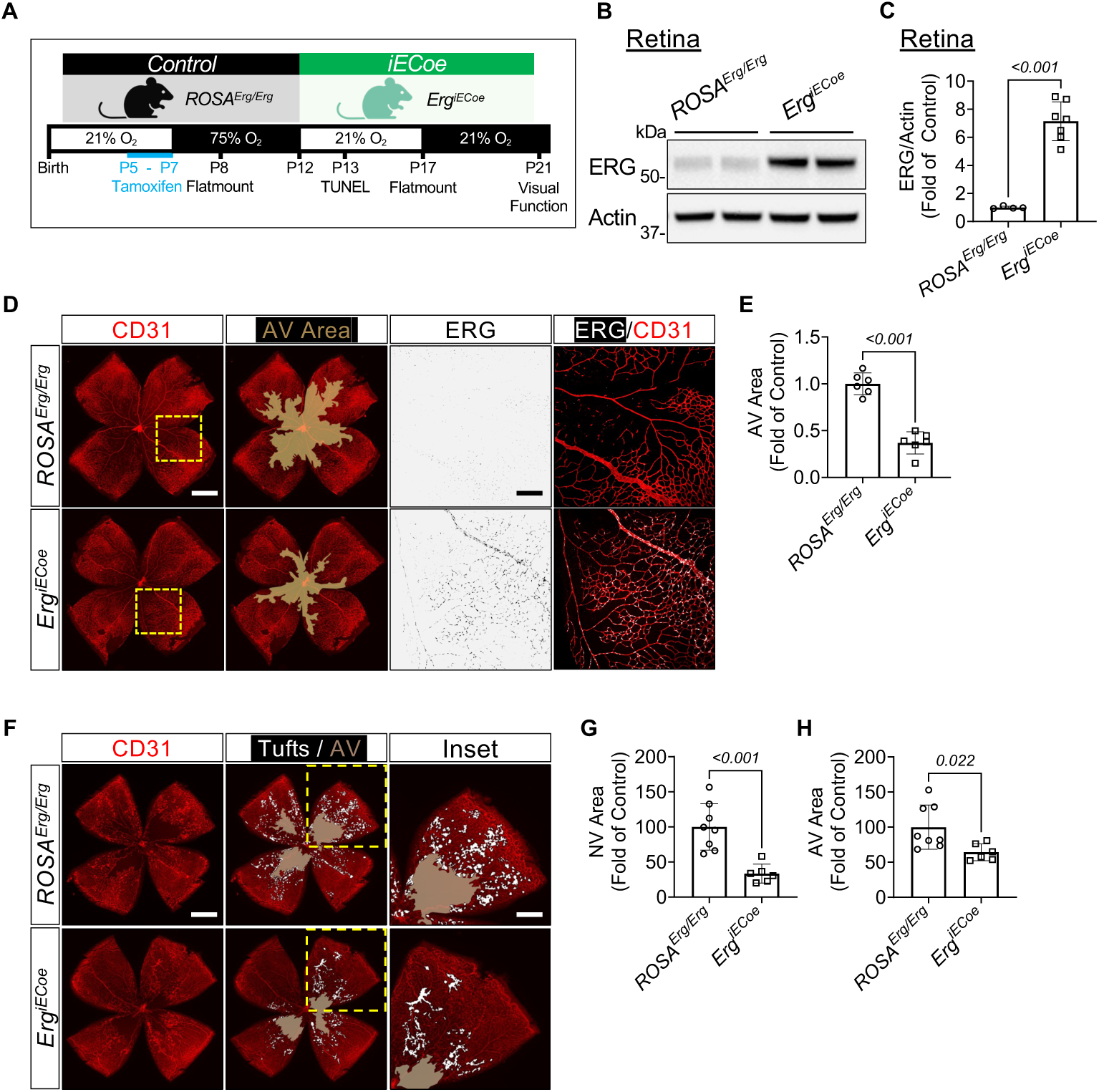
EC-specific ERG overexpression (*Erg^iECoe^*) reduces hyperoxia-induced regression and OIR-induced neovascularization. (**A**) Schematic of testing the effects of inducible endothelial ERG overexpression (*Erg^iECoe^*) in the OIR model. Tamoxifen was administered orally or by eyedrops to *Erg^iECoe^*and *Cre^ERT2^*-negative littermate control pups from P5 to P7. Vascular regression was measured at P8, and NV tufts were measured at P17. Neuronal damage was measured at P13 by TUNEL staining, and visual function was measured at P21 by electroretinograms and optokinetic tracking. (**B**) Representative immunoblots from whole retinas of P8 littermate controls and *Erg^iECoe^*pups after tamoxifen administration from P5 to P7. (**C**) Densitometry quantification of normalized ERG in (B) (n = 6-7 mice). (**D**) Representative immunostaining of CD31 (red) and ERG (white; insets) on retinal flatmounts at P8 after 24 hr of hyperoxia exposure. Avascular (AV) areas are labeled with beige layers. The inset magnified images correspond to the yellow squares in the images at the far left. Scale bar: 500 µm in the whole flatmount images and 150 µm in the magnified inset images. (**E**) Quantification of the central AV area in (D) (n = 6 mice). (**F**) Representative images of CD31-stained retinas from *Erg^iECoe^* pups and littermate controls at P17 after OIR challenge. SWIFT_NV was used to mark AV (beige) and NV tuft (white) areas. Scale bar: 500 µm in the whole flatmount images and 250 µm in the magnified insets. (**G, H**) Quantification of the neovascular (NV) areas (G) and AV areas (H) shown in (F) (n = 6-8 mice). Data are presented as mean ± SD. AV: Avascular, *Erg^iECoe^*: *ROSA^Erg/Erg^;Cdh5(PAC)-Cre^ERT2^*, NV: Neovascular.

After tamoxifen induction from P5 to P7 and exposure to hyperoxia for 24hr (Fig. 4A), immunofluorescence imaging of CD31-stained flatmounted retinas revealed that vascular regression was significantly attenuated in *Erg^iECoe^* mice compared to control littermates (Fig. 4D, 4E). We then asked if the observed reduction in hyperoxia-induced vascular regression led to a reduced neovascular response in *Erg^iECoe^* mice. At P17, the peak time of neovascularization in the OIR model, we observed reduction in both neovascular area and avascular area in *Erg^iECoe^*mice compared to littermate controls (Fig. 4F-H). This suggests that the prevention of early-stage vascular regression in the OIR model by ERG overexpression mitigates a compensatory neovascular response.

### Overexpression of endothelial ERG preserves retinal function following OIR

To determine if the preservation of retinal vessels in OIR-treated *Erg^iECoe^* mice had a positive impact on visual function, we induced endothelial ERG overexpression from P5 to P7, subjected mice to hyperoxia from P7 to P12, and then performed a TUNEL stain of retinal cross sections at P13 to assess neuronal cell death (Fig. 4A). We observed a significant reduction in OIR-induced apoptosis of neuronal retinal cells in *Erg^iECoe^*mice compared to littermate controls, particularly in the inner nuclear layer of the central retina (Fig. 5A, 5B).

**Figure 5.**
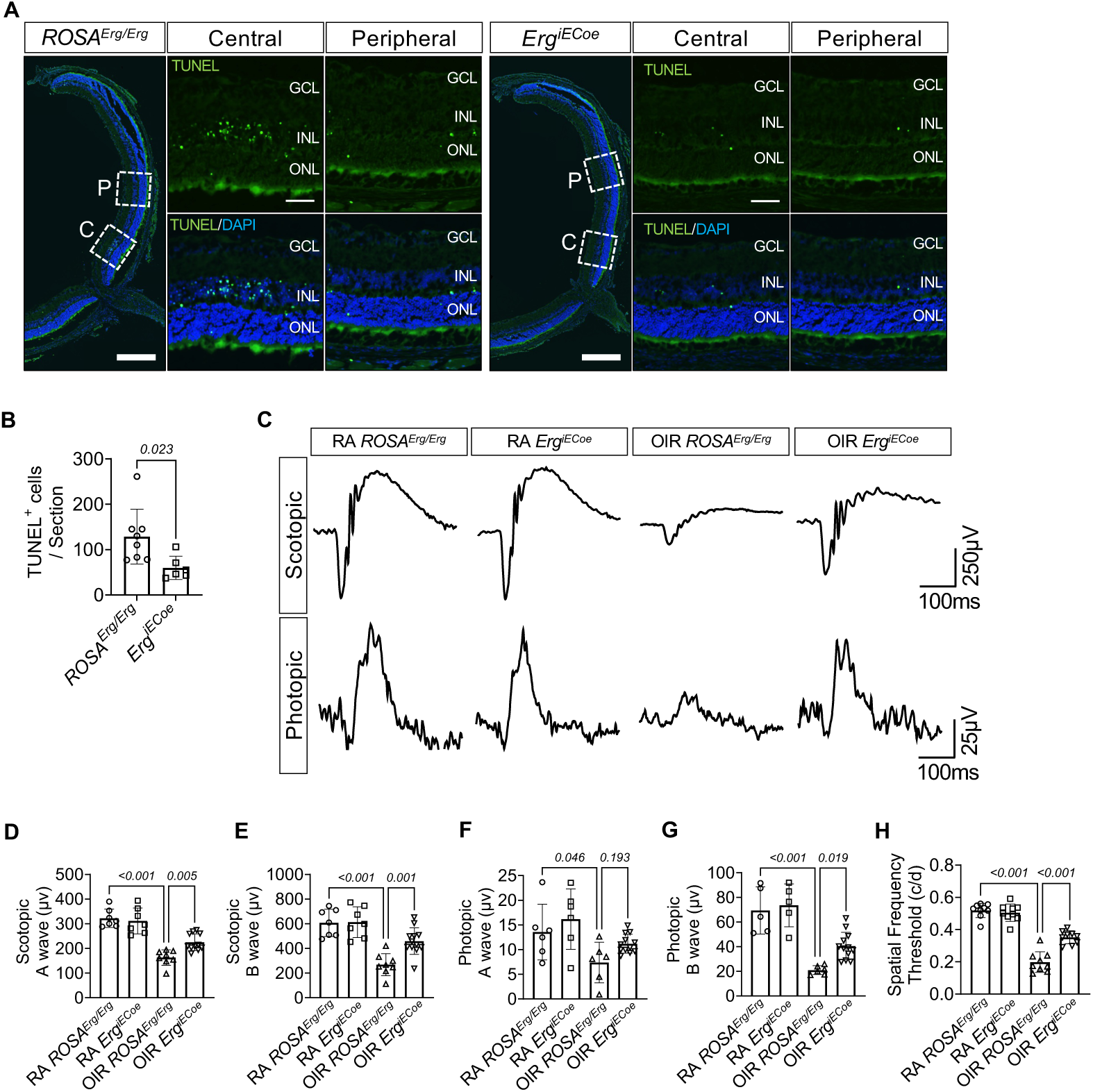
OIR-induced neural cell death and visual defects are reduced in *Erg^iECoe^* mice. **(A)** Following the OIR challenge schematic shown in Fig. 5A, TUNEL staining was performed on ocular cryosections of P13 *Erg^iECoe^*and littermate *Cre^ERT2^*-negative pups. DAPI was used as a counterstain. Scale bar: 250 µm in whole retinal sections and 25 µm in the magnified insets. (**B**) Quantification of TUNEL^+^ cells per whole retinal section (n = 6-8 mice). (**C**) Representative curves of scotopic and photopic amplitudes from both genotypes exposed to the OIR model or to room air (RA) as a negative control for the OIR model. Scale bar labels are shown at the bottom right. (**D-G**) Quantification of scotopic a-wave (D), scotopic b-wave (E), photopic a-wave (F), and photopic b-wave (G) readings as in (C) (n = 6-12 mice). (**H**) Quantification of spatial frequency threshold (c/d) by optokinetic tracking (OKT) (n = 8-9 mice). Data are presented as mean ± SD. C: Central, GCL: Ganglion cell layer, INL: Inner nuclear layer, OIR: Oxygen-induced retinopathy, ONL: Outer nuclear layer, P: Peripheral, RA: Room air.

The prevention of retinal neovascularization and the reduction of neuronal cell death in *Erg^iECoe^* mice prompted us to ask if these animals also had diminished OIR-induced visual defects. Therefore, *Erg^iECoe^*mice were administered tamoxifen from P5 to P7 (via eyedrop), exposed to 75% O_2_ (or room air) from P7 to P12, and then analyzed by electroretinogram at P21 (Fig. 4A). Note that both oral and eyedrop routes of tamoxifen administration resulted in comparably elevated ERG expression in *Erg^iECoe^* retinas (Suppl Fig. 4A and 4B), although *Erg^iECoe^* mice survived longer and appeared healthier at P21 when tamoxifen was administered via eyedrops. OIR-challenged *Erg^iECoe^* mice showed a significant improvement in a-and b-wave amplitudes for scotopic electroretinogram and b-wave amplitudes for photopic electroretinograms compared to control littermates (Fig. 5C-G). OIR-challenged *Erg^iECoe^* mice also showed a significant improvement in spatial frequency detection by OKT (Fig. 5H). Therefore, prevention of hyperoxia-induced retinal vascular regression correlates with improved visual function in the OIR model.

### ERG and FLI1 expression are downregulated in retinal capillaries of patients with early stages of DR

Finally, to determine the potential clinical significance of our findings, we assessed ERG and FLI1 expression in human eye sections from postmortem donors with non-proliferative DR (NPDR)—the initial stage of DR during which most retinal capillary regression occurs^2^. After immunostaining the eye sections, we found that both ERG and FLI1 were significantly downregulated in retinal capillaries of patients with NPDR compared to age-matched patients without DR (Fig. 6, yellow arrows). Notably, choroidal vessels, which are also visualized in retinal sections, do not regress in the early stage of diabetes and therefore can be used as an internal control ^36^. Importantly, we found that neither ERG nor FLI1 was downregulated in choroidal vessels of NPDR patients (Fig. 6, purple arrows), indicating that downregulation of ERG and FLI1 correlates specifically with ocular vessels undergoing regression at this stage of DR. Together with our OIR mouse studies, these results suggest that ERG and/or FLI1 downregulation is a conserved feature of early-stage retinopathies and a potential therapeutic target for mitigating disease progression.

**Figure 6.**
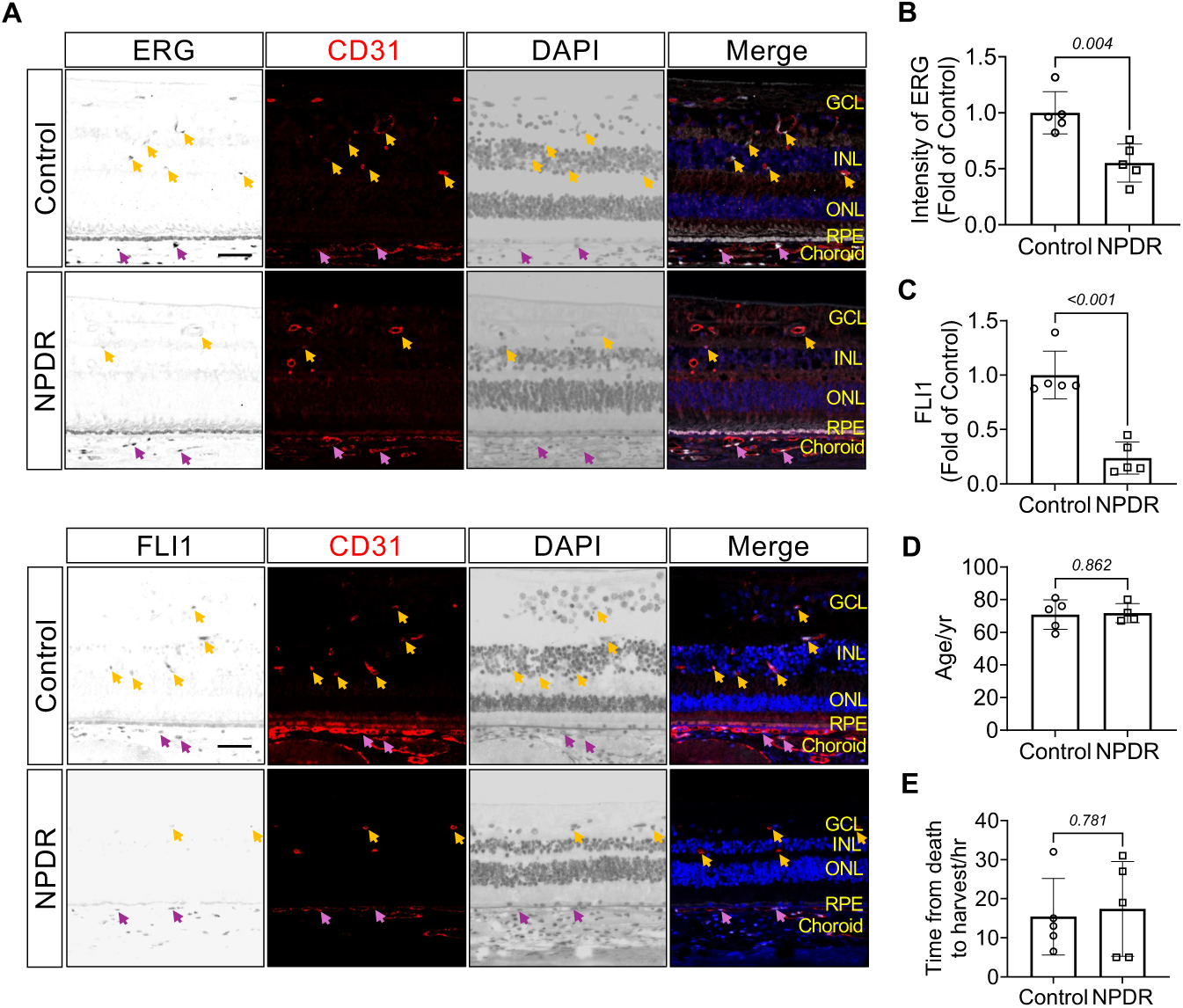
ERG and FLI1 are downregulated in retinal capillaries of patients with non-proliferative diabetic retinopathy (NPDR). Paraffin sections of postmortem human eyes were acquired from NPDR patients and age-matched individuals without DR (control). (**A**) Representative immunostaining of ERG, FLI1, CD31, and DAPI in human eye sections. Scale bar: 25 µm. Yellow arrows indicate retinal capillaries; Purple arrows indicate choroidal capillaries (CD31^+^DAPI^+^). (**B**) Quantification of ERG intensity in (A) (n = 5 individuals per group). (**C**) Quantification of FLI1 intensity in (A) (n = 5 individuals per group). (**D**) Ages of NPDR patients and age-matched controls. (**E**) The time from death to harvest of eyes from NPDR patients and age-matched controls. Data are presented as mean ± SD. GCL: Ganglion cell layer, INL: Inner nuclear layer, ONL: Outer nuclear layer, RPE: Retinal pigmented epithelium.

## DISCUSSION

Neovascular disorders of the eye are a substantial healthcare burden worldwide. There are ∼50,000 annual ROP diagnoses^7,37^, and incidence rates continue to rise due to improvements in survival rates for preterm infants^1^. In working-age populations, DR leads to blindness in ∼0.8 million individuals each year^38^, and in the United States more than 1.8 million patients (∼5% of all diabetes cases) are affected by vision-threatening DR^39^. Though there are substantial differences in the etiologies and affected populations of ROP and DR, both diseases are characterized by an initial loss of retinal vasculature that in turn leads to the formation of neovascular tufts. All currently available treatments target the stabilization of these neovascular tufts, and though they have efficacy in preventing severe consequences such as retinal detachment, no treatment yet addresses the underlying trigger of vascular insufficiency. This is due in part to a lack of understanding regarding the basic mechanisms of vascular regression that occur in ECs during the initiation of ROP and DR.

This study follows up on our previous work using a broad pharmacological inhibitor of ETS-family transcription factors to promote regression of neovascular tufts in the OIR challenge model^11^. Here, we use genetic models of EC-specific gene deletion to confirm that the loss of two specific ETS family members, *Erg* and *Fli1*, is sufficient to regress OIR-induced neovascular tufts. ERG and, to a lesser extent, FLI1 are predominantly expressed by ECs^17^. Because of the high homology of these proteins and previous observations of partially compensatory functions for ERG and FLI1^13,15^, it is not surprising that we needed to delete both *Erg* and *Fli1* to drive tuft regression. Although the identity of endothelial genes regulated by ERG and FLI1 during vascular regression was not addressed by this study, the homeostatic and stabilizing effects of these ETS factors on blood vessels are widely documented^12,13^. Nevertheless, the transcriptional targets of ERG and FLI1 can be contextually specific in vivo^13^, so future work will be needed to understand their transcriptional targets in retinal capillaries that are impacted by the OIR model.

Our data indicate that neovascular disorders of the eye offer two windows in which the regulation of vascular regression by ETS factors can be therapeutically useful. First, ERG and FLI1 inhibition can be used to promote selective regression of pathological neovascular tufts. This approach may offer a helpful complement to current interventions that focus on inhibiting VEGF to reduce both retinal neovascularization and vascular leakage that causes scarring and retinal detachment^3^. Secondly, stabilizing ERG expression can be used as a preventive measure to attenuate the hyperoxia-induced regression that initially drives tuft formation. Previous studies of hyperoxia-induced regression in the OIR model revealed that insufficient VEGF receptor signaling under hyperoxic conditions drives EC apoptosis^33,40,41^, and supplementation of VEGF or the alternative VEGF receptor ligand placental growth factor (PlGF) mitigates hyperoxia-induced retinal capillary regression^40,41^. Likewise, EC-specific deletion of the proapoptotic proteins BAK and BAX reduces this pathological regression process^33^. Our study adds to this body of literature by demonstrating the beneficial impact of preventing hyperoxic vessel regression on visual function. This indicates that photoreceptor damage is associated with early stages of the OIR challenge model, providing a possible explanation for OIR-induced visual defects that persist well after the spontaneous resolution of retinal neovessels^31^. Altogether, our study indicates that strategies preventing hyperoxic vessel regression may have additive benefits of alleviating both neuronal tissue ischemia and subsequent retinal neovascularization.

When we previously reported that pharmacological ETS factor inhibition could drive ocular vascular regression, we acknowledged that this process independently requires slow blood flow in capillaries^11^. Capillaries within neovascular tufts are thin and tortuous with slow flow as opposed to normal retinal vessels that maintain physiological flow, which explains why the broad ETS factor inhibitor that we administered to eyes of OIR-challenged mice at P17 could selectively regress neovascular tufts by P19. Hyperoxia drives retinal vasoconstriction in mice and humans^42,43^, indicating that both slow flow and ERG downregulation (Fig. 3) occur during the hyperoxia stage of the OIR model. DR is associated with more complicated retinal blood flow dynamics, although retinal capillaries are reported to have reduced diameters in early stages of this disease in humans and mice^44,45^. While future studies will be needed to address mechanisms through which slow flow and ETS factor downregulation collaboratively drive ocular vessel regression in early stages of neovascular eye diseases, our current study provides encouraging evidence that prevention of just one of these triggers—ETS factor downregulation—is sufficient to block the regression process.

Designing practical treatments to preserve ERG and FLI1 expression during the initiation of ROP and DR will require a better understanding of the mechanisms by which these ETS factors are downregulated during disease. Though ERG and FLI1 are highly expressed by ECs under normal circumstances, numerous studies have documented the downregulation of either factor during inflammatory diseases such as bacterial/viral infection, pulmonary arteriolar hypertension, autoimmune disease, liver disease, and aging^15,35,46–48^. Our new data raise the possibility that ERG expression is also regulated by direct oxygen exposure, which is consistent with published proposals that ERG expression is sensitive to hypoxia^15^ and to oxidative damage by H_2_O_2_^49^. Interestingly, we previously reported that ERG has heightened protein turnover in ECs of the lung^35^, an organ notable for its exposure to environmental oxygen levels. Therefore, we speculate that a similar proteolytic mechanism may result in oxygen-dependent ERG downregulation in blood vessels of the eye. Specifically, our data raise the possibility that oxygen-induced ERG reduction is due to O_2_-dependent regulation of the ubiquitin ligase TRIM25, which has previously been shown to mediate ERG proteolysis in ECs^34^. TRIM25 function has largely been characterized in non-ECs where it participates in viral DNA responses, although a recent study has shown TRIM25 to play a role in ECs during oxidative stress^50^. Interestingly, this study focused on TRIM25 function under hyperglycemic conditions, suggesting that TRIM25-mediated ERG degradation may be common to both ROP and DR. Still, future studies will be needed to explore the mechanisms by which oxygen—or other metabolic stimuli in the context of DR—promotes ERG degradation.

## NOVELTY AND SIGNIFICANCE

What is known?

- Ischemic retinopathies are characterized by initial vascular regression that leads to subsequent pathological neovascularization.
- Some erythroblast transformation-specific (ETS) transcription factors promote endothelial cell (EC) homeostasis and stability.
- A broad pharmacological inhibitor of ETS factors promotes the regression of neovascular tufts in the murine oxygen-induced retinopathy (OIR) model that mimics retinopathy of prematurity (ROP).

What new information does this article contribute?

- EC-specific deletion of the ETS factors ERG and FLI1 promotes neovascular tuft regression in OIR-challenged mice, although this regression does not improve visual function, which is damaged in earlier stages of the OIR model.
- Endothelial ERG expression is downregulated in early stages of the OIR model, and EC-specific overexpression of ERG reduces the initial OIR-induced capillary regression that drives subsequent photoreceptor death, neovascularization, and vision impairment.
- Patients with early stages of diabetic retinopathy (DR) exhibit lower ERG and FLI1 expression in retinal capillaries compared to people without diabetes, indicating that loss of these ETS factors may be a conserved contributor to capillary regression that initiates retinopathies.

This study highlights the importance of ERG and FLI1 transcription factors in regulating vascular regression at different stages of ischemic retinopathies. Using EC-specific gene deletion, we found that loss of these ETS factors promoted regression of OIR-induced pathological neovascular vessels in the eye but failed to improve visual function, suggesting that relevant retinal damage occurs prior to and independently of neovascularization. Turning our attention to the earliest stage of the OIR model, we found reduced ERG expression in retinal ECs of wild type mice, raising the possibility that oxygen-induced ERG downregulation promotes vessel regression during the initiation of OIR-induced pathology. We therefore developed a murine model of EC-specific ERG overexpression and found it sufficient to prevent hyperoxia-induced vascular regression, neuronal cell death, and neovascularization in the OIR model. Importantly, ERG overexpression also improved visual function in OIR-challenged mice, demonstrating that inhibition of early-stage capillary regression blocks subsequent pathological sequelae associated with ROP. Finally, we show that both ERG and FLI1 are downregulated in retinal vessels of human patients with early stages of diabetic retinopathy, suggesting that neovascular disorders of the eye may share common mechanisms underlying pathological retinal capillary regression.

## ACKNOWLEDGEMENTS

The authors appreciated advice about immunostaining the human eye sections from Dr. Ching Yuan and Ms. Heidi Roehrich (Lions Gift of Sight Eye Bank, University of Minnesota). The authors are also grateful for histological support from Dr. Wenjing Wu (Dean McGee Eye Institute) and training in visual function tests from Drs. Ana Chucair-Elliott, Willard Freeman, and Scott Plafker (Oklahoma Medical Research Foundation).

## SOURCES OF FUNDING

This work was supported by grants from the NIH (R35HL144605, P20GM139763), the British Heart Foundation (RG/11/17/29256 and RG/17/4/32662) the American Heart Association (23SCEFIA1155941 and 24POST1196957), and an EyeFind Research Grant from the ARVO Foundation for Eye Research.

## DISCLOSURES

CTG, CMS, and EM filed a provisional patent (#63/666,397) related to this work.

## AUTHOR CONTRIBUTIONS

Conceptualization: EM, CMS, CTG

Methodology: EM, CMS, AMR, GMB, CTG

Investigation: EM, CMS, JX, YR, JK, CTG

Visualization: EM, CMS, CTG

Funding acquisition: EM, CMS, AMR, CTG

Project administration: CTG

Supervision: CTG

Writing – original draft: EM, CMS, CTG

Writing – review & editing: EM, CMS, AMR, GMB, CTG

## DATA AND MATERIALS AVAILABILITY

All data are available in the main text or the supplementary materials. The following mice were obtained and used through MTAs: *Fli1^flox^* (Boston University) and *Cdh5(PAC)-Cre^ERT2^* (Cancer Research Technology Limited). *ROSA^Erg^* mice are available through Imperial College London with MTA.

## Notes

### Competing Interest Statement

The authors have declared no competing interest.

