## Supplemental Figs 1-4 and Table 1 for "Targeting endothelial ERG to mitigate vascular regression and neuronal ischemia in retinopathies"

**Affiliations:**

### Supplementary Figures

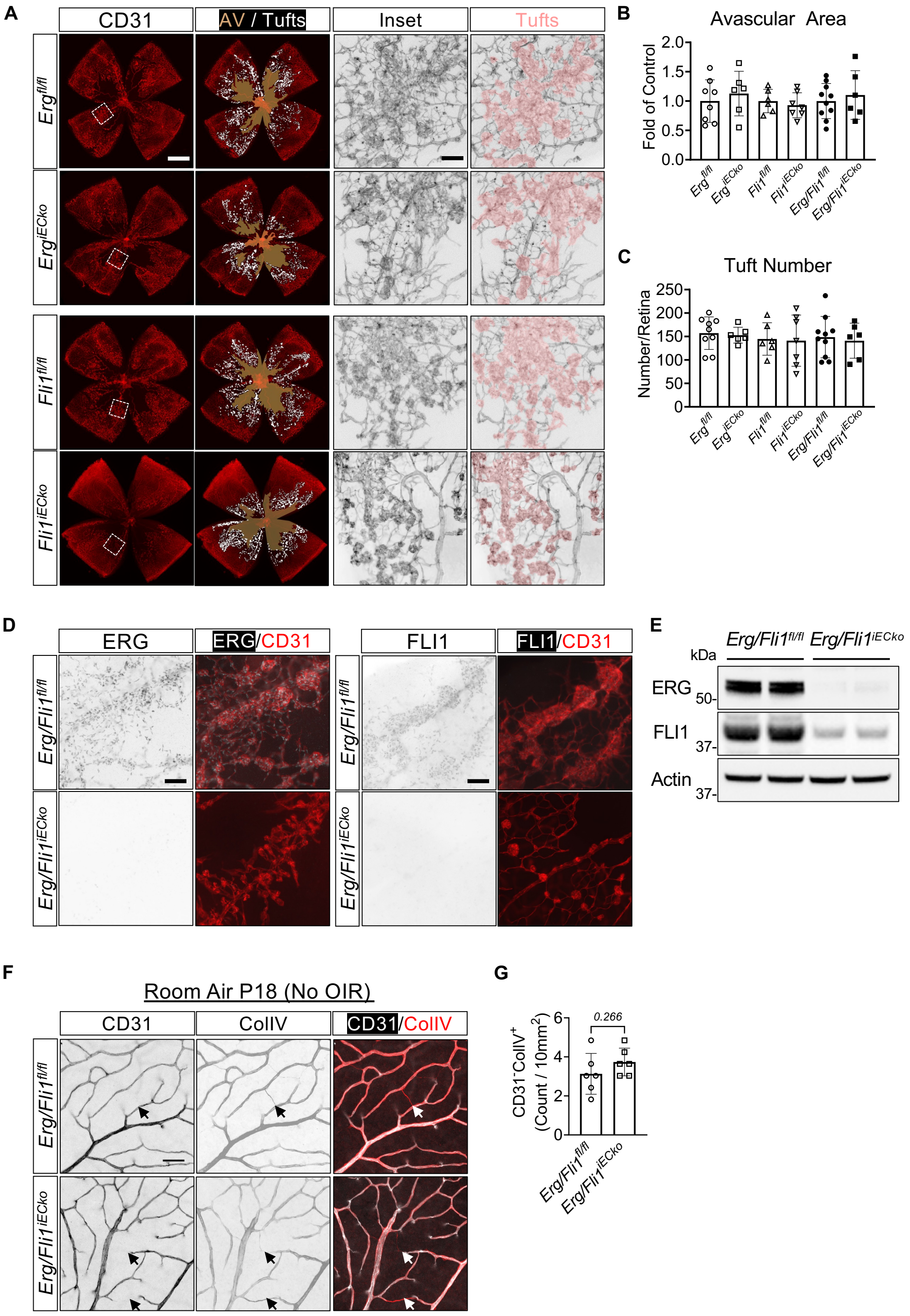


**Supplementary Figure 1. EC-specific deletion of either *Erg* or *Fli1* does not promote NV tuft regression.**

(**A**) Representative images of CD31-stained retinal flatmounts from P18 *Erg*$^{fl/fl}$ and *Erg*$^{iECko}$ mice, and littermate control mice. The ImageJ plugin SWIFT_NV (*17*) was used to label and quantify NV areas (white in the whole flatmounts) and avascular (AV) areas (beige layer). The inset monochrome images correspond to the white squares in the images at the far left; tufts are pseudocolored in pink. Scale bar: 500 µm in whole flatmounts and 50 µm in the magnified inset images. (**B**) Quantification of AV area from *Erg*$^{iECko}$, *Fli1*$^{iECko}$, *Erg/Fli1*$^{iECko}$, and littermate control mice (n = 6-10 mice). (**C**) Quantification of total tuft number from *Erg*$^{iECko}$, *Fli1*$^{iECko}$, *Erg/Fli1*$^{iECko}$, and littermate control mice (n = 6-10 mice). (**D**) Representative images of ERG, FLI1, and CD31 staining from P18 *Erg/Fli1*$^{iECko}$ and control mice after tamoxifen administration and OIR challenge, as shown in Fig. 1A. Scale bar: 50 µm. (**E**) Representative immunoblots of ERG and FLI1 in retinal lysates from P18 *Erg/Fli1*$^{iECko}$ and control mice after tamoxifen administration and OIR challenge, as shown in Fig. 1A. (**F**) Representative images of empty basement membrane sleeves (arrows; CD31$^{-}$ColIV$^{+}$) from P18 *Erg/Fli1*$^{iECko}$ mice and littermate control mice that were raised in room air. Scale bar is 50 µm. (**G**) Quantification of empty basement membrane sleeves in (F) (n = 6 mice). Data are presented as mean ± SD. AV: Avascular; OIR: Oxygen-induced retinopathy.

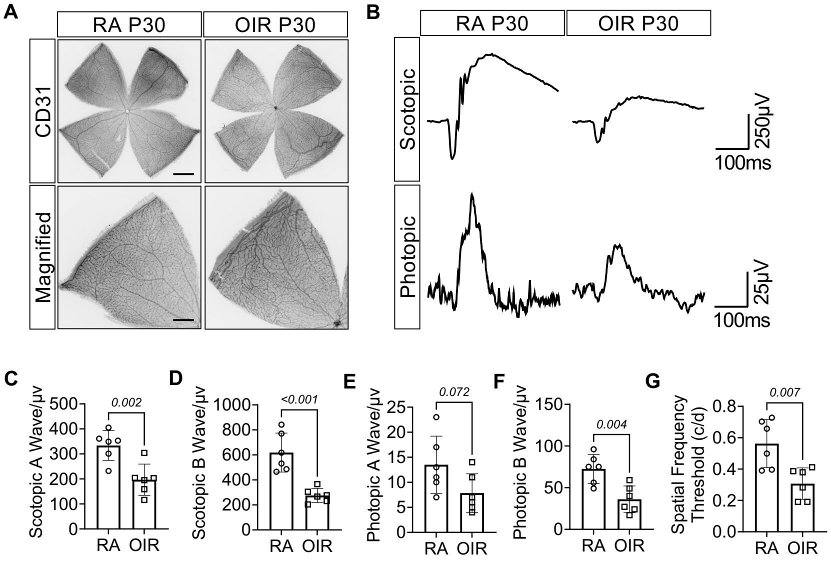


**Supplementary Figure 2. Visual function is still compromised in wild type mice at P30 after OIR challenge.**

WT C57BL/6J mice were challenged with the OIR model. Electroretinogram and optokinetic tracking (OKT) were measured at P30, and retinas were subsequently harvested for whole flatmount immunostaining. (**A**) Representative images of CD31-stained retinal vasculature from littermate mice challenged with OIR or raised in room air (RA) as controls. Scale bar: 500 µm in whole flatmounts and 200 µm in the magnified images. (**B**) Representative curves of scotopic and photopic amplitudes from P30 OIR and RA mice. Scale bar labels are shown at the bottom right. (**C-F**) Quantification of scotopic a-wave (C), scotopic b-wave (D), photopic a-wave (E), and photopic b-wave (F) in (B) (n = 6 mice). (**G**) Quantification of spatial frequency threshold (c/d) measured by OKT (n = 6 mice). Data are presented as mean ± SD. OIR: Oxygen-induced retinopathy, OKT: Optokinetic tracking, RA: Room air.

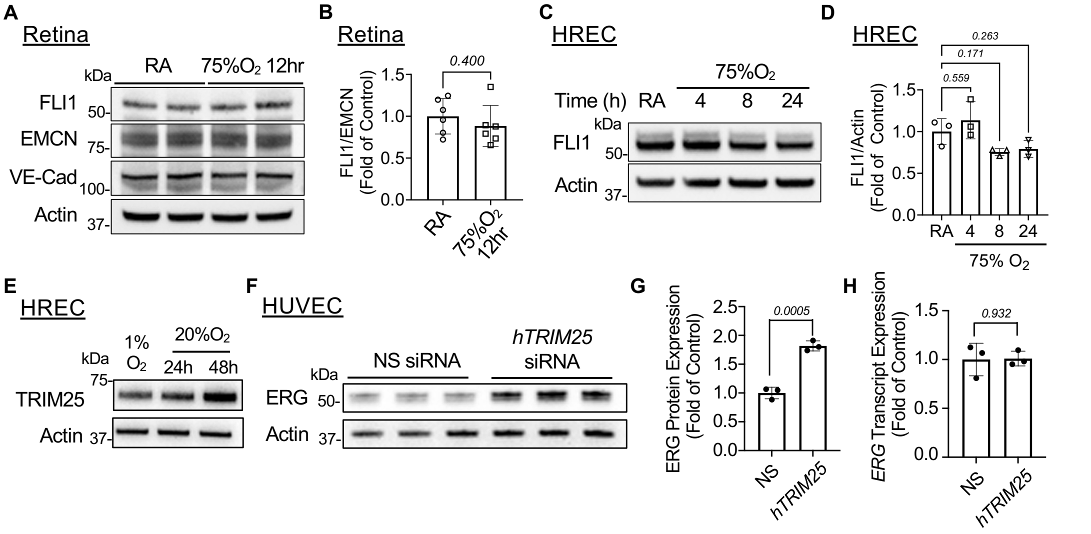


**Supplementary Figure 3. Hyperoxia potentially degrades ERG but not FLI1 via upregulation of the ubiquitin ligase TRIM25.**

(**A**) Representative immunoblots from whole retinas of P7 littermate wild type pups exposed to room air (RA) or to 75% $O_2$ for 12 hr. (**B**) Densitometry quantification of FLI1 normalized to endomucin (EMCN) (n = 6 mice). (**C**) Representative immunoblots from primary human retinal microvascular endothelial cells (HREC) cultured under RA for 24 hr or 75% $O_2$ for the indicated time. (**D**) Densitometry quantification of normalized FLI1 in (C) (n = 3 technical replicates). (**E**) Immunoblot for TRIM25 protein expression in HRECs cultured under 1% $O_2$ or 20% $O_2$ for 24 or 48h. (**F-H**) Primary Human Umbilical Vein ECs (HUVECs) were treated with non-specific (NS) or *hTRIM25*-targeting siRNAs for 72h followed by quantification of ERG protein and transcript expression by immunoblot (F and G) and qPCR (H), respectively (n = 3 technical replicates). Data are presented as mean ± SD. EMCN: Endomucin, HREC: Human retinal microvascular endothelial cells, HUVEC: Human umbilical vein endothelial cells, RA: Room air, VE-Cad, Vascular endothelial cadherin.

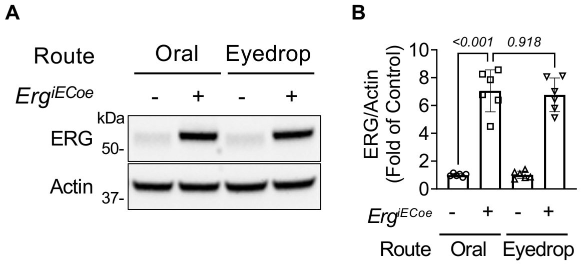


**Supplementary Figure 4. ERG overexpression is comparable in *Erg^iECoe^* pups after oral or eyedrop routes of tamoxifen administration.**

**(**A) Representative immunoblots from whole retinas of P8 *Erg*$^{iECoe}$ pups and littermate controls. Tamoxifen was administered via oral or eyedrop routes from P5 to P7. (**B**) Densitometry quantification of ERG normalized to Actin in (A) (n = 6 mice). Data are presented as mean ± SD.

**Supplementary Table 1. List of antibodies used in this study.**

| Antibodies | Host | Company | Catalog No. | Application | Dilution |
|---|---|---|---|---|---|
| CD31 | Goat | R&D | AF2836 | IF | 1:100 |
| CD31 | Rat | BD Biosciences | 553370 | IF | 1:100 |
| Collagen IV | Goat | Millipore | AB769 | IF | 1:100 |
| Endomucin | Rat | Santa Cruz | sc-65495 | WB | 1:500 |
| ERG | Rabbit | Abcam | ab92513 | 1) Western blotting (WB); 2) Immunofluorescence (IF) | 1) 1:2000; 2) 1:100 |
| FLI1 | Rabbit | Abcam | ab15289 | 1) WB; 2) IF | 1) 1:1000; 2) 1:100 |
| TRIM25 | Rabbit | Abcam | ab167154 | WB | 1:4000 |
| VE-Cadherin | Mouse | Santa Cruz | sc-9989 | WB | 1:500 |
| β-Actin | Mouse | Santa Cruz | sc-47778 | WB | 1:2000 |
| Anti-goat IgG | Donkey | Vector Laboratories | PI-9500 | WB | 1:3000 |
| Anti-mouse IgG | Horse | Vector Laboratories | PI-2000 | WB | 1:3000 |
| Anti-rabbit IgG | Goat | Vector Laboratories | PI-1000 | WB | 1:3000 |
| Anti-goat IgG (Alexa Fluor 647) | Donkey | Jackson ImmunoResearch Laboratories Inc. | 705-607-003 | IF | 1:250 |
| Anti-rabbit IgG (Alexa Fluor 594) | Donkey | Jackson ImmunoResearch Laboratories Inc. | 111-585-003 | IF | 1:250 |
| Anti-rat IgG (Alexa Fluor 594) | Donkey | Jackson ImmunoResearch Laboratories Inc. | 712-585-153 | IF | 1:250 |
